# Evidence Accumulates for Individual Attributes during Value-Based Decisions

**DOI:** 10.1101/2021.08.05.455296

**Authors:** Douglas G. Lee, Todd A. Hare

**Affiliations:** Institute of Cognitive Sciences and Technologies, National Research Council of Italy; Zurich Center for Neuroeconomics, Department of Economics, University of Zurich; Neuroscience Center Zurich, University of Zurich and ETH Zurich

**Keywords:** multi-attribute choice, value-based choice, preferential choice, sequential sampling model, drift-diffusion model

## Abstract

When choosing between different options, we tend to consider specific attribute qualities rather than deliberating over some general sense of the options’ overall values. The importance of each attribute together with its quality will determine our preference rankings over the available alternatives. Here, we test the hypothesis that the most prominent class of model for simple decisions – sequential sampling or evidence accumulation to bound – can be bolstered by explicitly incorporating variables related to individual attributes in addition to the standard usage of overall value estimates. We examine six datasets in which participants evaluated snack foods both in terms of overall value and individual attributes, then chose between pairs of the same snacks, and show that only models that explicitly incorporate information about the individual attributes are able to reproduce fundamental patterns in the choice data, such as the influence of attribute disparity on decisions, and such models provide quantitatively better fits to the choice outcomes, response times, and confidence ratings compared to models based on overall value alone. Our results provide important evidence that incorporating attribute-level information into computational models helps us to better understand the cognitive processes involved in value-based decision- making.

## Introduction

Most decisions that we make are based on information about a variety of relevant features of the available options. Theories and mathematical models of multi-attribute choice generally agree that, *in principle*, the decision system in our brains should compare options based on how well they score cumulatively across all relevant attribute dimensions (Bettman et al., 1998; E. J. Johnson & Payne, 1985; Keeney et al., 1993; Levav et al., 2010; Payne et al., 1988, 1993; Russo et al., 1996; Shah & Oppenheimer, 2008). These overall scores, be they based on subjective valuations or more objective features, are typically thought to be calculated as the weighted sums of sub-scores across all dimensions (Bettman et al., 1998; Bhatia & Stewart, 2018; E. J. Johnson & Payne, 1985; Levav et al., 2010; Payne et al., 1988, 1993; Russo et al., 1996; Shah & Oppenheimer, 2008). Specifically, each option will be assigned a score along each attribute dimension, and each dimension will be given some weight according to how relevant or important it is to the decision. Simplifications of this strategy that assign equal weights to all attributes (Dawes, 1979; Dawes & Corrigan, 1974), reduce attribute scores to binary better/worse rankings (Russo & Dosher, 1983), or only consider a subset of the attributes have also been proposed (Fishburn, 1974; Tversky, 1972). How well these simpler strategies perform depends on the choice context (Bettman et al., 1998; Gigerenzer & Gaissmaier, 2011; E. J. Johnson & Payne, 1985; Levav et al., 2010; Payne et al., 1988, 1993; Russo et al., 1996; Shah & Oppenheimer, 2008). Regardless of precisely how they are combined, almost every choice is determined by an assessment of multiple attributes. Thus, it is important for both basic and applied researchers to better understand how the attribute composition of choice options (and not just their overall values) influences the decision-making process in the brain.

Recent work has investigated the question of whether value-based decisions are driven by attribute-level comparison processes, and specifically whether such processes might lead to choice behavior that systematically varies as a function of the disparity of the options’ attribute compositions (Lee & Holyoak, 2021). A pair of options has high disparity if, for example, one option scores high in the first attribute dimension but low in the second, while the other option scores high in the second dimension but low in the first (Figure 1a, left panel). On the contrary, a pair of options has low disparity if both options have similar scores along each attribute dimension (Figure 1a, middle panel). Notably, two decisions could be equally difficult in the traditional sense that the overall value ratings of the choice options are equally close together, yet have very different levels of disparity (Figure 1a, right panel). In multiple independent experiments, Lee & Holyoak (Lee & Holyoak, 2021) found that choice behavior differs as a function of disparity, such that higher disparity corresponds to higher choice consistency (i.e., a choice in favor of the option that was previously rated as having the higher overall value), lower response time, and higher choice confidence, even after accounting for differences in overall values. Here, we re-examine the Lee & Holyoak (Lee & Holyoak, 2021) data using computational modeling and show that models of value comparison that assume an immutable combination of attributes into the overall option value cannot account for this pattern of results.

**Figure 1.**
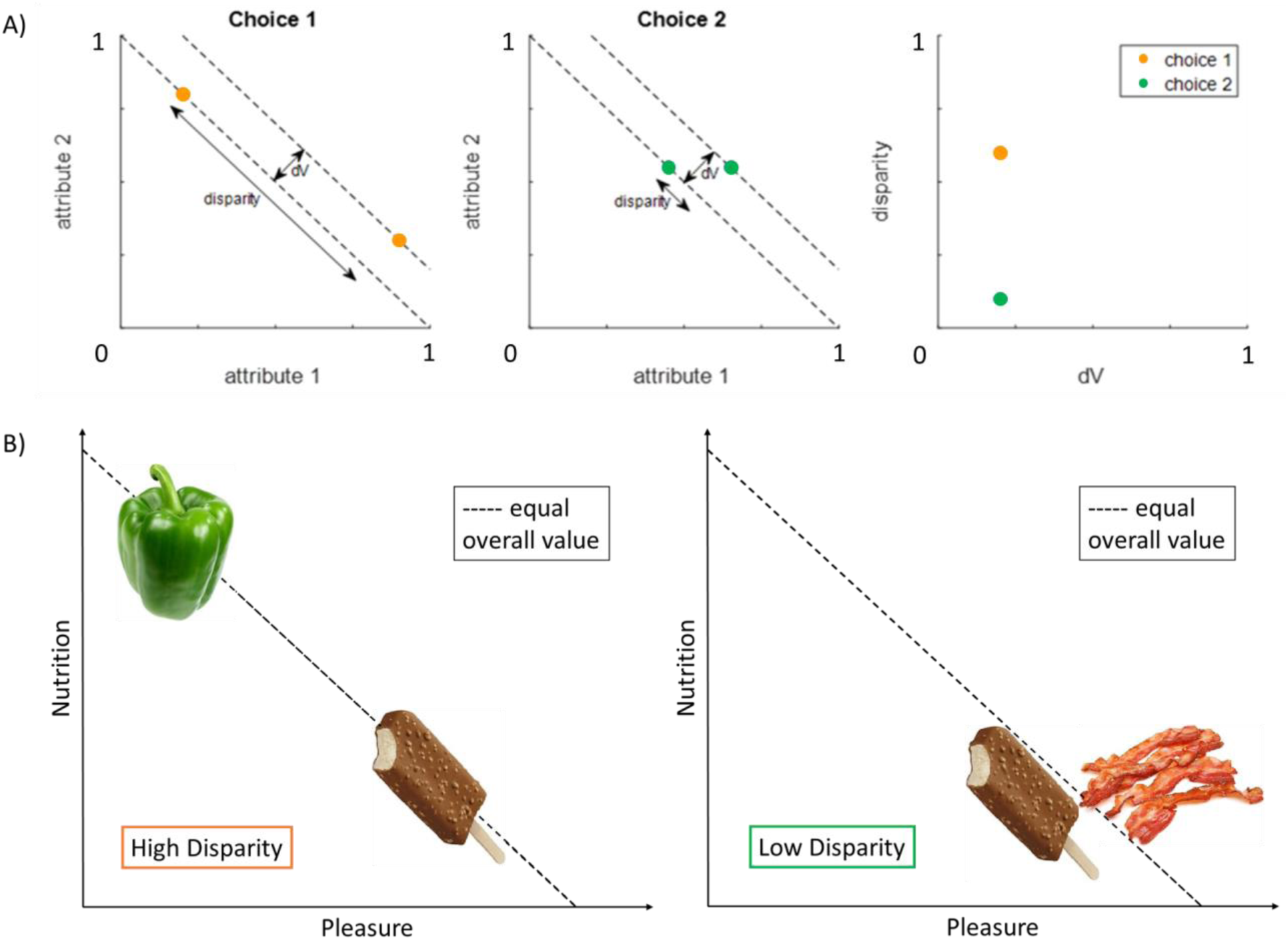
Choice disparity. **a)** A schematic illustration of orthogonal components of choice difficulty: dV and disparity. The left plot illustrates a “high disparity” choice, and the middle plot illustrates a “low disparity” choice. The orange and green dots represent the alternative options for each choice, each plotted according to its measurements on two attribute dimensions. The example assumes equal importance weights for each attribute, so the iso-value curves are represented by parallel lines with slope -1. The difference in overall value of the options, dV, is the distance between the iso-value curves on which the options lie. Disparity is the distance between the options in the dimension orthogonal to overall value (see Equation 1 below for a mathematical formulation). The right plot shows the location of each choice pair in the transformed dV-disparity space. **b)** An example illustration of two choice sets for snack foods, one high disparity (left plot), one low disparity (right plot). As shown by the dashed iso-value lines, all of the available snacks are of comparable overall value (and thus each choice pair is of comparable low dV). However, the two choice pairs are of very different disparity. In the high disparity pair (left), one option scores high on pleasure but low on nutrition, while the other option scores low on pleasure but high on nutrition. In the low disparity pair (right), both options score high on pleasure and low on nutrition.

Instead, choices between naturalistic multi-attribute stimuli and subsequent confidence reports for those choices are best explained by models in which individual attributes are actively (re)weighted during the comparison process. In our tests, we focus on multi-attribute choices between naturalistic unitary options with multiple inherent latent features, as opposed to bundled or conjoint options made up of multiple components (e.g., probability + amount for risky choice; delay + amount for inter-temporal choice; effort or pain + reward for cost-benefit tradeoffs; different items for bundled choices). We believe that this type of naturalistic reward, which could plausibly be treated as an integrated whole, provides a stronger test of whether items are compared based on fixed weights that yield stable overall values or values constructed from flexible attribute weights during decisions. We find that models that allow context-dependent attribute weights during decisions best explain the outcome, response time, and confidence data. This demonstrates not only that attributes are considered individually, but also that the manner in which attributes contribute to value estimates (measured either explicitly through ratings or implicitly through choices) might vary across context (rating versus choice). However, we also show that using a subset of attribute-specific ratings together with overall value ratings can help to better explain the choice data (e.g., when obtaining ratings for the full set of potentially relevant attributes is impractical).

In the domain of risky choice, it has long been established that individuals’ preferences over options that combine monetary gains and losses with probabilities or time delays may reverse when different methods are used to elicit those preferences. For example, preferences revealed through choices have been shown to reverse compared to those elicited by matching, pricing, or rating procedures (Alós-Ferrer et al., 2016; Alos-Ferrer et al., 2020; Alós-Ferrer et al., 2021; Fischer et al., 1999; Grether & Plott, 1979; Lichtenstein & Slovic, 1971; Seidl, 2002; Tversky et al., 1988, 1990; Weber & Johnson, 2009). A leading explanation for these preference reversals is that the importance weights on the probability, time, and/or money dimensions differ across the preference elicitation procedures (Seidl, 2002; Tversky et al., 1988). Eye-tracking experiments have shown that changes in the proportion of visual fixations to the monetary amount of a lottery’s potential outcome relative to its probability across choice and pricing trials are associated with the differences in the relative weights given to amounts versus probabilities when choosing versus setting a price (Alós-Ferrer et al., 2021). This influence of visual attention on context-dependent weighting is consistent with sequential sampling models that predict that the effects of overall value and attribute differences on choices are determined in part by the amount of attention paid to each option or attribute (Busemeyer & Townsend, 1993; Diederich, 1997; Krajbich et al., 2010; Roe et al., 2001). Together, these theories and data form the basis of our hypothesis that decision values in naturalistic multi-attribute choices will also be constructed at the time of choice from the options’ basic attributes in a context-dependent manner, rather than being compared as unitary overall values aggregated across all attributes in a constant fashion. We aim to show that individual attributes directly influence the valuation process in consumer choices over unitary goods whose attributes must be recalled from memory rather than read off a screen. We propose that attribute-level evidence accumulation is necessary and sufficient to explain the observed disparity effects on choice (Lee & Holyoak, 2021).

## Methods

### Data

We analyzed the data from six previous experiments ((Lee & Holyoak, 2021); Experiments 1-5 plus an unpublished pilot experiment that we label Experiment 0). Experiments 1-5 recruited a total of 325 participants (186 female, mean age = 40, age range: 19-50) and Experiment 0 recruited 55 participants (demographic data were not recorded for this unpublished experiment). All participants were native English speakers from the United States or Canada, recruited via Amazon Mechanical Turk. The total number of participants included in our analyses was 307 (41 for Experiment 0, 50 for experiment 1, 48 for experiment 2, 54 for experiment 3, 60 for experiment 4, and 54 for experiment 5), using the same exclusion criteria from the original study: using Spearman correlation coefficients between first and last ratings (within participant) in addition to slope parameters from logistic regression of choice on value (separately using first and last ratings), participants with scores outside a cutoff (median +/− 3 × median average deviation) were excluded.

In each experiment, participants completed several distinct phases. They first passively observed images of individual snack foods (100 in Experiments 0 and 1, 60 in Experiments 2-5). Second, they provided overall value ratings for each individual snack food. While we and the original authors refer to these as ratings of overall value, the participants were never actually directly asked about value. Rather, they were asked to respond to the question, “How much would you like this as a daily snack?” Participants entered their response to this question for each snack, separately, before proceeding to the next experimental section. Next, they rated the pleasure they expected to derive from each food and its nutritional value, in separate experimental sections (the order of the pleasure and nutrition rating tasks was counterbalanced across participants). In Experiments 2-5, participants then completed a “non-choice” task (e.g., stating how similar paired options were to each other), followed by another series of rating tasks (overall value, pleasure, and nutrition) identical in format to the initial rating tasks. In all experiments, participants then completed a choice task in which they chose their preferred snack from pairs of options (50 choice trials for Experiments 0 and 1, 30 choice trials for Experiments 2-5). During the choice section, after each choice, participants also rated their confidence that the option they chose was indeed their preferred option on that trial. Lastly, participants completed a final series of rating tasks (overall value, pleasure, and nutrition) identical in format to the previous rating tasks. For the purposes of our current study, we focus on those ratings that occurred immediately prior to the choice task (i.e., the first series of ratings for Experiments 0-1 and the second series of ratings for Experiments 2-5). We also examine the post-choice ratings, to rule out the possibility that the reported results only hold for pre-choice ratings. We did not expect to find any differences by rating timing, since the pre- and post-choice ratings were highly correlated (cross-participant median Pearson correlation coefficient for overall value, pleasure, and nutrition, respectively: 0.90, 0.90, 0.95). The consistency of the pre- and post-choice ratings indicates that participants likely reported their true opinions during each of the rating sessions. This is because random ratings would not be as consistent, and maintaining intentionally altered ratings of attributes or overall value relative to one another would require exerting substantial cognitive effort for no real benefit, which is something that humans rarely do. See Figure 2 for an illustration of the different tasks.

**Figure 2.**
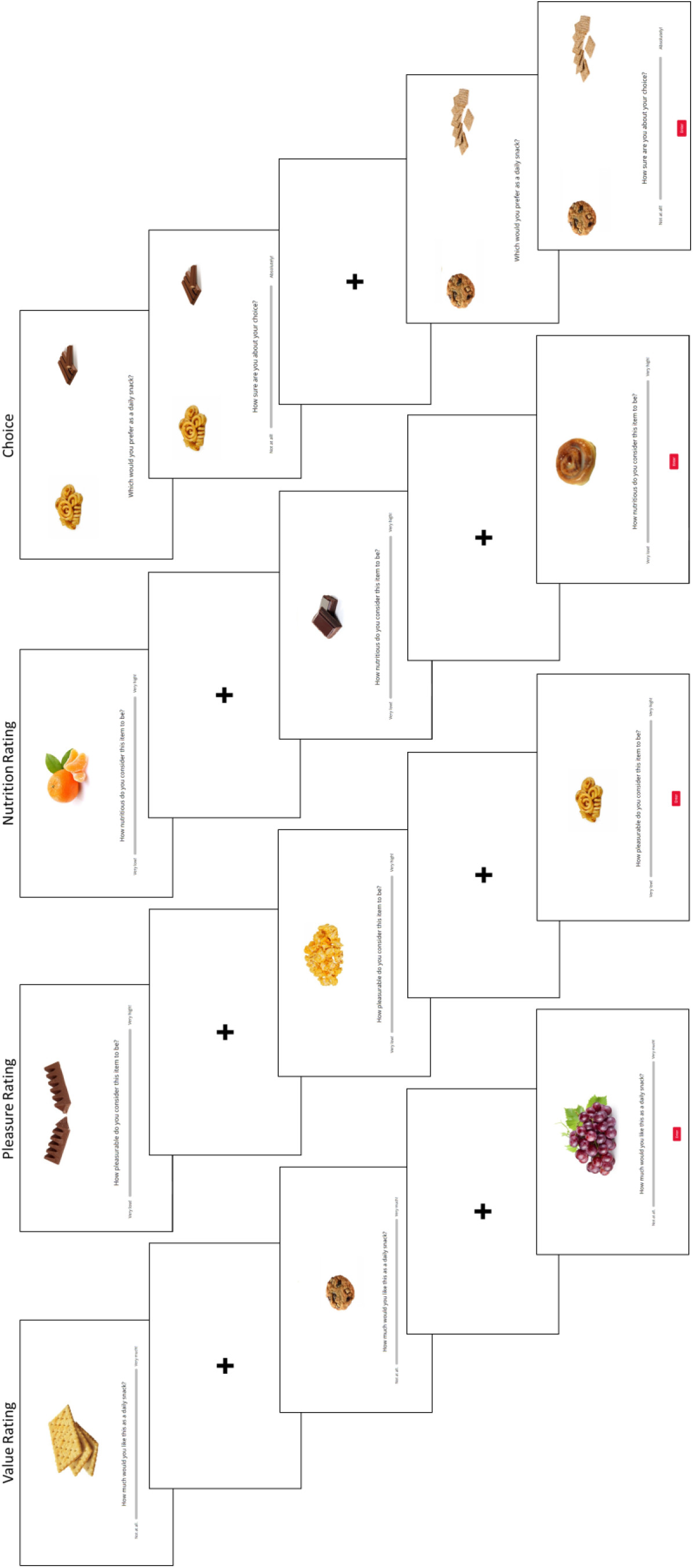
Task illustration. An example of what participants saw on the screen during each experimental section.

In addition to the aforementioned datasets, we also examined data from an unpublished pilot auxiliary task that were originally collected along with the primary data reported in (Lee & Daunizeau, 2021). The participants in this study were native French speakers, recruited via the RISC platform at the Paris Brain Institute (demographic data not available). In this study, all participants provided overall value ratings for 148 food options, then made 74 choices between pairs of options, then again provided overall value ratings for the same 148 options. Out of the main group of participants, 17 completed the auxiliary task (determined solely by which participants had extra time remaining in their assigned testing time slot), in which they rated each of the food options in terms of “taste”, “health”, “texture”, and “appearance”. The format of the different tasks was similar to that described above, except that all text was presented in the French language. We are thus able to compare versions of our models (described below) that incorporate four attributes rather than two (the third and fourth attributes enter the models in the same way as the first and second attributes).

### Evidence accumulation models

In this study, we consider several ways in which attribute and overall value estimates might influence the evidence accumulation process in a sequential sampling model (Bhatia, 2013; Busemeyer et al., 2019; Pleskac & Busemeyer, 2010; Ratcliff, 1978; Ratcliff & McKoon, 2008; Ratcliff & Rouder, 1998; Roe et al., 2001; Trueblood et al., 2014; Usher & Mcclelland, 2001). The general class of sequential sampling models can include various features and assumptions such as inhibition, leakage, non-linear drift rates, or time-varying drift rates. While different sequential sampling models vary in their predictions about the exact shape of the choice response time distributions, they share the assumption that decision accuracy or consistency and mean response times are proportional to drift or evidence accumulation rates (faster evidence accumulation leads to faster responses and more correct/consistent choices). Because our aim is to show how evidence accumulation rates relate to ratings of overall value and individual attributes, without making any claims about other potential model complexities, we examine variants of a basic drift-diffusion model (DDM; (Ratcliff, 1978; Ratcliff & McKoon, 2008; Ratcliff & Rouder, 1998)). In its simplest form, the DDM includes a drift (evidence accumulation) rate, a measure of diffusion noise, and a measure of separation between response thresholds triggering a choice of either option. The specifics of each model that we examined are detailed below. Under the DDM, the values of the options (in simple two-alternative forced-choice tasks) are repeatedly compared across time. The so-called evidence that arises in favor of one option over the other is corrupted by processing (e.g., neural) noise, so repeated samples are accumulated to cancel out the noise. Once a sufficient amount of evidence has been accrued (i.e., the response threshold is reached), the process terminates and a choice is made. The various types of evidence accumulation model that operate over differences in evidence for each option or allow inhibition between separate accumulators all make qualitatively similar predictions about the relationship between the evidence accumulation rate and choice probabilities, response times (RTs), and confidence ratings. Specifically, greater evidence accumulation rates are associated with more correct or consistent choices, faster RTs, and higher levels of confidence. We tested the fit of these generalized predictions to data when the evidence accumulation rate is proportional to the difference in overall value between the options (OV) or to a weighted linear combination of the differences in individual attributes (IA) using six different empirical data sets. We demonstrate that the evidence accumulation based on a weighted linear combination of the differences in individual attributes provides a better account of choice probabilities, mean RTs, and the effects of attribute disparity on choice outcomes and RT.

### Evidence accumulation based on overall value (OV)

In this case, evidence about the overall value of each option is sampled at each time step, the evidence for the two options is compared, and the relative evidence in favor of option 1 over option 2 is aggregated by the evidence accumulator. For simplicity, we use a basic framework based on a minimal version of the drift diffusion model (DDM, (Ratcliff, 1978; Ratcliff & McKoon, 2008; Ratcliff & Rouder, 1998)) with symmetric and constant boundaries to derive quantitative predictions about choices, mean response times, and confidence levels. Within our basic DDM framework, we assume that the cumulative evidence (*x*) evolves across deliberation time as follows:

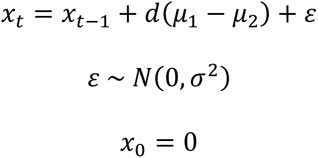

where *d* is an efficiency parameter for the evidence accumulation (the drift rate in a DDM), μi is the reported overall value of option *i* ∊ {1, 2}, and σ^2^ is the strength of the white noise ε in the accumulation process. Evidence sampling and accumulation proceeds until *x* reaches a response boundary ∊ {θ, -θ}, with the sign determining the chosen option (arbitrarily defined as positive for option 1, negative for option 2). Response time (RT) is equal to *t* at the moment a boundary is crossed. General predictions about the choice probability (*p*, choice of option 1) and mean RT from a basic DDM can be analytically derived (Alós-Ferrer, 2018) as a function of μ1, μ2, and σ^2^, with θ being fixed (here, to θ = 1 for simplicity):

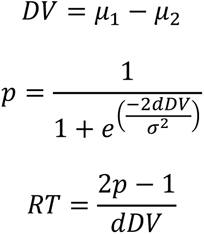

where μ1 and μ2 are independent variables, and *d* and σ^2^ are free parameters to be estimated to capture the individual-specific mean rate of evidence accumulation (drift rate) and level of noise in the accumulation process, respectively. We focus on mean response times because those are more similar across the different flavors of evidence accumulation models than the entire response time distribution, and our aim is to test general predictions independent of any specific model.

### Evidence accumulation based on individual attributes (IA)

In this case, evidence about the individual attributes of each option is sampled at each time step, the evidence for the two options is compared separately for each attribute, and the relative evidence in favor of option 1 over option 2 for each attribute is aggregated by the evidence accumulator. The IA model is of the same form as the OV model, except that the evidence accumulator is driven by two evidence streams (one for each attribute dimension: a, b). The process is otherwise identical, and it unfolds as follows:

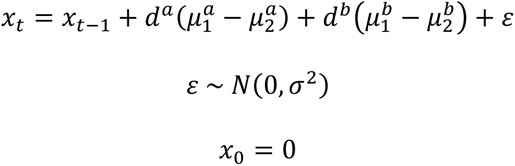

where μi^j^ is the reported value of option i ∊ {1,2} along attribute dimension j ∊ {a,b}, and σ^2^ is white noise common to the overall evidence accumulation process. Choice probability and mean RT are derived as:

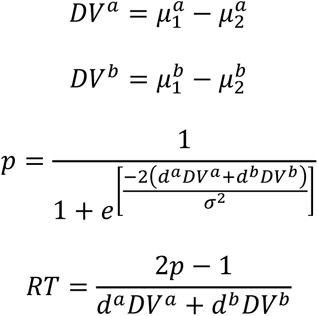

where d^j^ are independent free parameters that allow for different rates of evidence accumulation within each attribute dimension j∊ {a,b}.

### Evidence accumulation based on overall value and individual attributes (IA+)

In this case, evidence about both the overall value and the individual attributes of each option is sampled at each time step, the evidence (for both OV and IA) for the two options is compared, and the relative evidence in favor of option 1 over option 2 is aggregated by the evidence accumulator. This IA+ model is of the same basic form as the OV and IA models. However, the IA+ assumes that the drift rate is driven by the separate values of the individual attributes that were explicitly evaluated (in these experiments, pleasure and nutrition), but that it is also influenced by other attributes that were not explicitly evaluated. Thus, if the overall value ratings contain information about the attributes that were rated individually as well as other attributes that were not rated, including overall value should enhance the model fit. Therefore, in this model, the evidence accumulator is driven by evidence streams for each explicit attribute dimension as well as the aggregate overall value estimates. The process is otherwise identical to that in the OV and IA models, and it unfolds as follows:

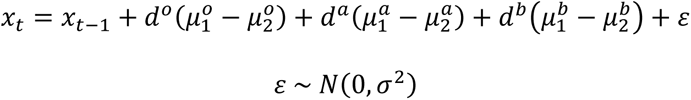

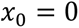

where μi° is the reported overall value for option *i* ∊ {1, 2}, μi^j^ is the reported value of option i ∊ {1,2} along attribute dimension j ∊ {a,b}, and σ^2^ is white noise common to the overall evidence accumulation process. (Note that in cases where the individual attributes are highly correlated and/or the attribute ratings jointly explain a large portion of the variance in overall values, it may be necessary to employ orthogonalization or dimensionality reduction techniques, if the goal is to make inferences about the relative weights or importance of attributes in determining the drift rate.) Choice probability and mean RT are derived as:

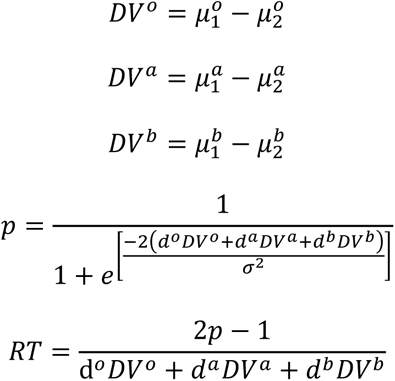

where d^j^ are independent free parameters that allow for different rates of evidence accumulation for overall value estimates and within each attribute dimension j∊ {a,b}.

### Model fitting procedure

We fit each of the three candidate models to the experimental data, then performed Bayesian model comparison to determine which of the models (if any) performed significantly better than the others across the population of participants. For this model fitting and comparison exercise, we relied on the Variational Bayesian Analysis toolbox (VBA, available freely at https://mbb-team.github.io/VBA-toolbox/; (Daunizeau et al., 2014)) with Matlab R2020a. Within participant and across trials, we entered the experimental variables {ratings of overall value, pleasure, and nutrition for each option} as input and {choice = 1 for left option, 0 for right option; RT; choice confidence} as output. All ratings and reports of certainty and confidence were rescaled to range from (0-1]. We also provided the model-specific mappings from input to output as outlined in the analytical formulas above. As we fixed the threshold parameter θ to 1, the parameters to be fitted were thus the drift rate *d* and diffusion noise σ^2^ terms described above in the model formulations. VBA requires prior estimates for the free parameters, for which we set the mean equal to 1 and the variance equal to 23 for each parameter. The theoretical drift rate and noise parameters are always positive; we thus constrained the search space of our model fitting algorithm to the positive domain by replacing these parameters with the following calculation: log(1 + exp(parameter)) * 2.3^1^. VBA then recovers an approximation to both the posterior density on unknown variables and the model evidence (which is used for model comparison). We used the VBA_NLStateSpaceModel function to fit the data for each participant individually, followed by the VBA_groupBMC function to compare the results of the model fitting across models for the full group of participants.

One benefit of using VBA to fit the data to our models is that it is computationally efficient, as it relies on Variational Bayesian analysis under the Laplace approximation. This iterative algorithm provides a free-energy approximation for the model evidence, which represents a natural trade-off between model accuracy (goodness of fit, or log likelihood) and complexity (degrees of freedom, or KL divergence between priors and fitted parameter estimates; see (Friston et al., 2007; Penny, 2012)). This is a critical step for comparing models that differ in number of parameters.

Additionally, the algorithm provides an estimate of the posterior density over the model’s free parameters, estimated from our flat Gaussian priors. Individual log model evidence scores are then provided as input to the group-level random-effect Bayesian model selection (BMS) procedure. BMS provides an exceedance probability that measures how likely it is that a given model is more frequently implemented, relative to all other models under consideration, in the population from which participants were drawn (Rigoux et al., 2014; Stephan et al., 2009). This approach to fitting and comparing variants of DDM has already been successfully demonstrated in previous studies (Feltgen & Daunizeau, 2021; Lee & Usher, 2021; Lopez-Persem et al., 2016). Our VBA-based approach makes use of the concise analytical formulation of mean RT, as opposed to the full distribution of RT. To verify that this simplification did not bias model fits or comparisons, we repeated the model comparison after fitting the data using the full RT distributions and choice probabilities (but without choice confidence) with the R package runjags (Denwood, 2016; R Core Team, 2020) as an interface for JAGS (Plummer, 2003; Wabersich & Vandekerckhove, 2014). The model comparison results were the same as what we report below (see Supplementary Material).

An additional benefit of fitting and comparing models within the VBA toolbox is that it allows us to fit the formulas for choice probability and mean RT in a generic manner by simultaneously optimizing parameters over a collection of analytical equations. Moreover, while choice probability and RT are summarized by a set of two equations, we can just as easily perform our model comparison while fitting additional equations alongside those for choices and response times. Therefore we fit generalized predictions about the relationship between the evidence accumulation rate and choice confidence in addition to choice probability and mean RT. We estimated choice confidence as a linear function of either overall value difference (OV), individual attribute differences (IA), or both (IA+). In each case, we assume that higher drift rates would be associated with greater confidence. This is consistent with empirical data showing that confidence decreases monotonically with choice difficulty (which determines the drift rate), as well as previous studies investigating the relationship between evidence accumulation and confidence (Pleskac & Busemeyer, 2010). The formulas that we used to estimate confidence under the different models were thus:

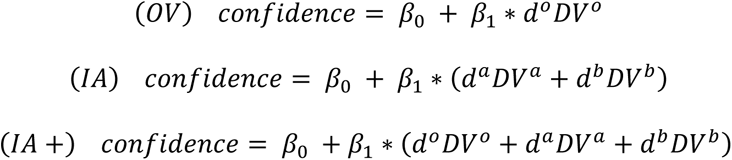

### Model recoverability

To verify that our model-fitting procedure is suitable for this specific analysis, we performed a test of model recoverability. Specifically, we created a group of 1,000 synthetic participants, each with 50 choice trials. For each trial, we generated a random number in the range: (0-1) for each of the pleasure and nutrition ratings. For each synthetic participant, we generated a random relative weight parameter (RW) for the importance of pleasure versus nutrition, in the range: (0-10). We then generated the value of each option by sampling from a normal distribution with mean equal to the weighted average of the pleasure and nutrition ratings and random standard deviation in the range: (0,1). We assigned random parameters for each participant under each model by sampling from normal distributions with the following means and standard deviations: drift rates: (1, 0.3) for dV under OV, (0.2, 0.3) for dN under IA, (0.5, 0.3) for dV under IA+, (0.1, 0.2) for dN under the IA+; diffusion noise: (0.5, 0.1) under all models; confidence: (0.5, 0.2) for both parameters under all models. For IA and IA+, the drift rate parameters for dP were set equal to the drift rate for dN * RW. We then simulated the set of choice probabilities, mean RTs, and confidence ratings for each participant, separately according to each of our models. Finally, we fit all simulated data (per synthetic participant) to each of our models and performed the same formal model comparison as with our real experimental data. The results of this procedure can be seen in Figure 3A as a model confusion matrix. This matrix shows, for each true generative model, the percentage of synthetic participants (under that model) that were attributed to each of the best fit models by our model-fitting procedure. As shown in the matrix, model confusion was low and the procedure attributed the true generating model as the best fitting model for the vast majority of the simulated participants (recovery accuracy: 99% for OV, 86% for IA, 85% for IA+).

**Figure 3.**
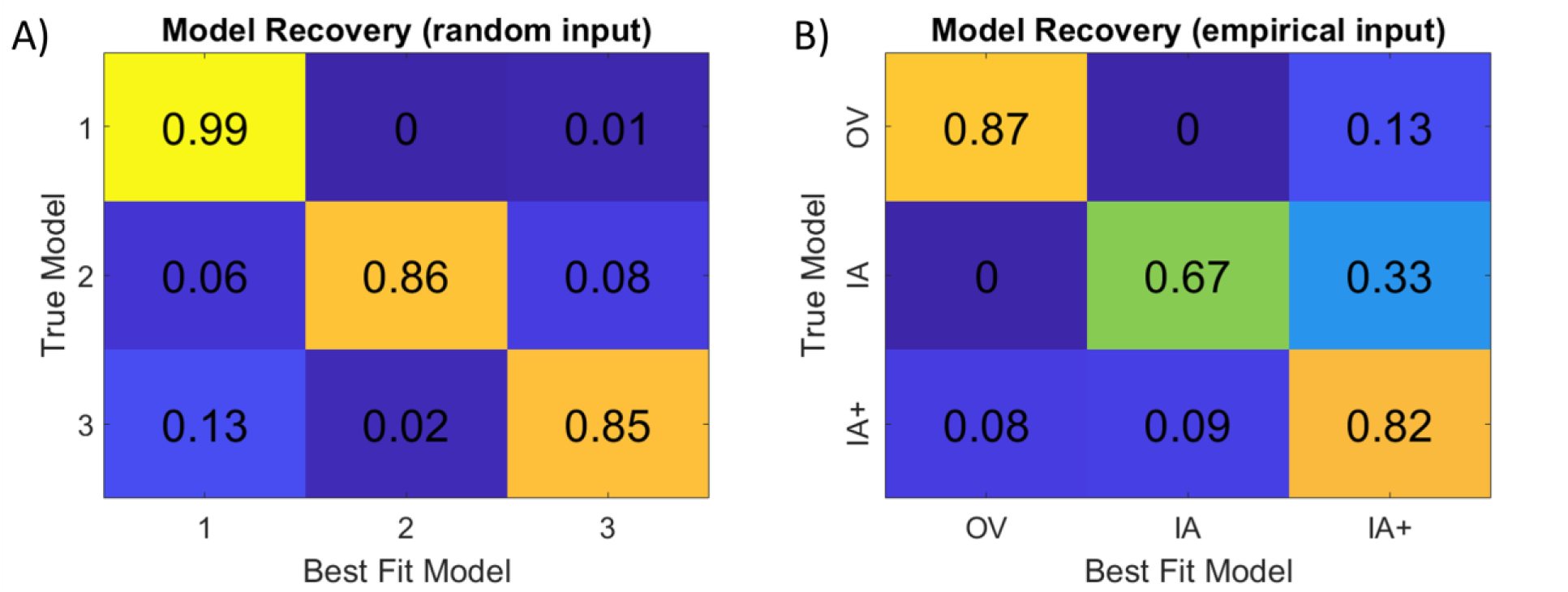
Model recovery analysis using random input (A) and empirical input (B). The cells in each “confusion matrix” summarize the percentage of simulated participants (under each true model) for which our model-fitting procedure attributed each of the (best fit) models.

We then needed to determine if the model recoverability would remain high when using the precise distributions of variables observed in our empirical data. We thus repeated the model recovery analysis based on real data. Specifically, we took as the model input the actual data for each participant in Studies 1-6 (ratings of overall value, pleasure, and nutrition for each option). We then simulated the set of choice probabilities, mean RTs, and confidence ratings for each participant, separately according to each of our models, using the actual participant-specific fitted parameters for each model. Finally, we fit all simulated data (per participant) to each of our models and performed the same formal model comparison as with our real experimental data. The results of this procedure can be seen in Figure 3B. As shown in the matrix, model confusion remained very low (recovery accuracy: 87% for OV, 67% for IA, 82% for IA+). There was still some confusion between the IA+ and other specifications, which is expected because the IA+ is essentially a combination of those other specifications.

## Results

### Relationship between variables

Our main analyses rely on some fundamental assumptions about how the different variables (value, pleasure, and nutrition) relate to each other. First, it is necessary that the pleasure and nutrition attributes explain a large portion of the variance in overall value ratings, i.e., that those attributes are an adequate subset of the relevant attributes for these choices. We tested this by running a linear regression of value on pleasure and nutrition, separately for each participant. Across participants, the GLM beta weights for pleasure and nutrition were 0.736 and 0.192, respectively (*p* < .001 for both, based on 2-sided t-tests of each beta versus zero), and the mean r- squared value for the multiple regressions was 0.730. Second, it is necessary that pleasure and nutrition are sufficiently orthogonal and explain unique variance in overall value ratings and choices. We tested this by calculating the Pearson’s correlation between pleasure and nutrition ratings, separately for each participant. Across participants, there was no systematic correlation between pleasure and nutrition (mean ρ = -0.034, *p* = .137, based on a 2-sided t-test versus zero).

There was heterogeneity across participants, but most correlation coefficients were within the [- 0.53, 0.52] range (80% of participants; see the Supplementary Material).

### Qualitative Model Predictions

The evidence accumulation models we compare make distinguishable predictions regarding the impact of two important features of binary choices: overall value difference and attribute disparity. Lee and Holyoak (Lee & Holyoak, 2021) introduced the term *disparity* as a secondary characteristic of a decision, orthogonal to the primary characteristic most often used in the field: overall value difference (often referred to as choice difficulty, where lower value difference implies higher difficulty). The disparity between two options {i,j} with respect to two attribute dimensions (here, P for pleasure and N for nutrition) is calculated as:

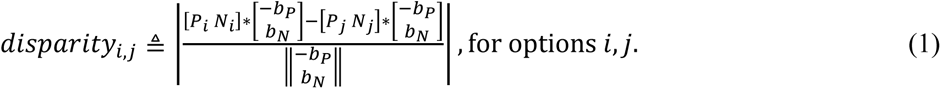

where bP and bN are the importance weights for the P and N attributes, respectively. Note that it would be straightforward to extend this formula to incorporate more than two options or more than two attributes. Disparity calculated in this way effectively transforms the variable space from attribute dimensions to decision dimensions (see Figure 1 above).

We first show the qualitative predictions that each evidence accumulation process (OV, IA, IA+, all simulated under their participant-specific best fitting parameters) makes with respect to the effects of value difference (dV) and disparity (D) on choice consistency, RT, and choice confidence, and how this compares to the empirical data (Figure 4). To generate the synthetic data, we used the real trial-by-trial values faced by each individual participant (value of option 1, or v1; value of option 2, or v2; pleasure of option 1, or p1; pleasure of option 2, or p2; nutrition of option 1, or n1; nutrition of option 2, or n2), along with each participant’s best-fitting parameters for each of the three evidence accumulation specifications. Note: we coded the data such that option 1 always had the higher overall value. Next, we performed mixed effects (MFX) regressions of choice (binomial), of RT (linear), and of confidence (linear) on dV and D, pooling all simulations together and including study and participant as random effect regressors. The IA+ specification is the only one from the set we examined that is able to account for the relative magnitudes and directionality of the associations dV and D show with choice consistency (whether or not the higher-rated option was chosen), RT, and confidence (see Figure 4). Note that because the simulated data are less noisy than the empirical data, those regression coefficients will be larger (i.e., the regressors will explain more of the variance in the dependent variables). Regardless, what is important here is the relative magnitudes of the coefficients for dV and D *within* each data source, not the absolute magnitudes or comparisons across data sources.

**Figure 4.**
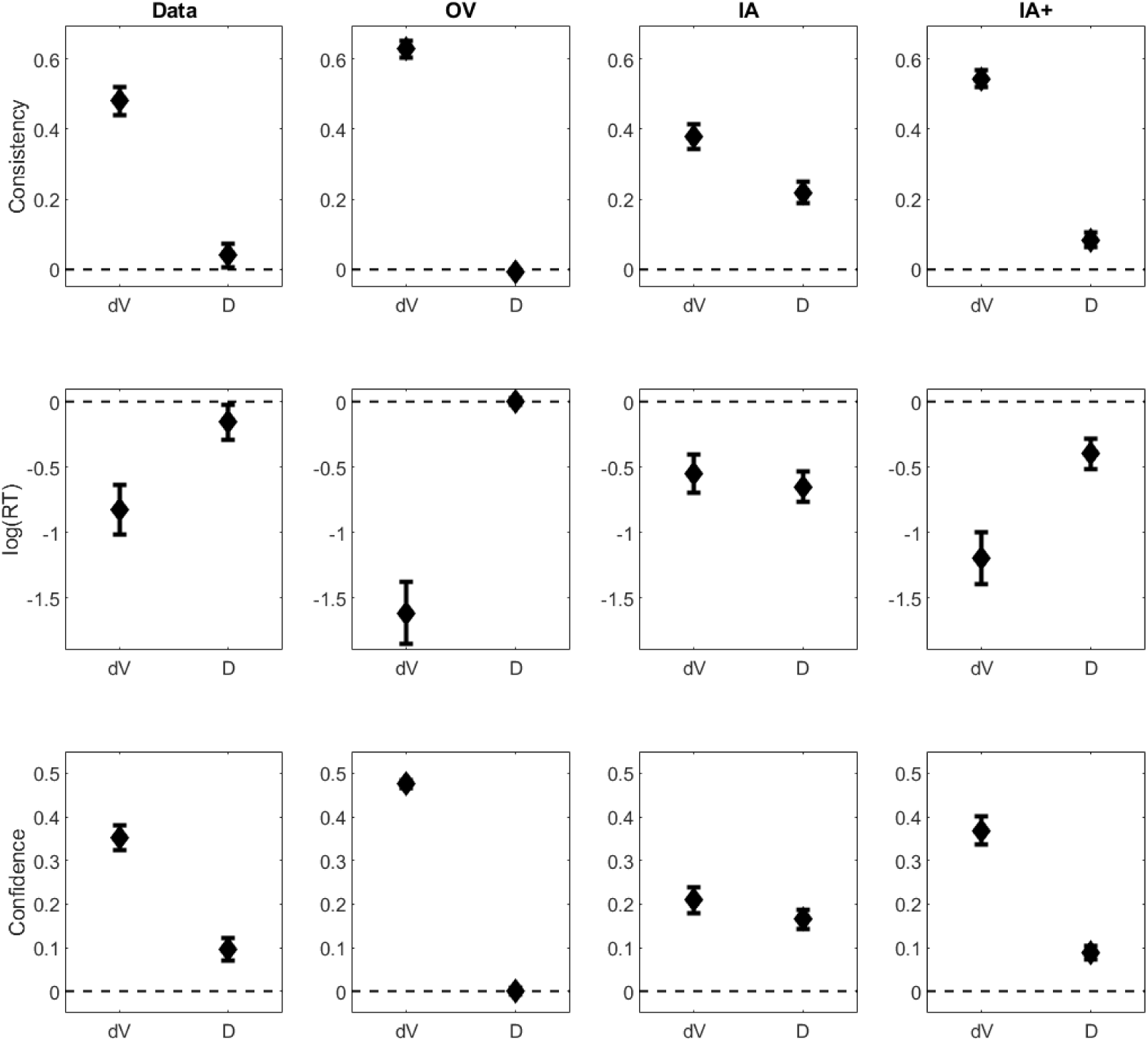
Model predictions. Qualitative predictions of the effects of value difference (dV = value of option 1 – value of option 2) and disparity (D; see equation 1) on choice consistency, log(RT), and confidence in the empirical (column 1) and simulated data (columns 2-4; shown for responses simulated using the best fitting parameters for each model); diamonds represent MFX fixed effects regression coefficients; error bars represent 95% confidence intervals.

### Quantitative model comparisons

To quantitatively evaluate the models, we performed formal quantitative model comparisons of the various diffusion decision model specifications. We compared the fit of the model based on overall value (OV) to a model based on individual attribute values (IA), to test our hypothesis that decision makers place different importance or decision weights on attributes during choices than when rating the overall value of each option in isolation. If the attributes are weighted and aggregated the same way during ratings and choices, then the OV model, which uses the comprehensive overall value of the foods to estimate choices should outperform the IA model, which only uses a subset of the foods’ attributes (pleasure and nutrition) to estimate choice outcomes, response times, and reported confidence levels. If instead attributes are weighted differently during choices compared to ratings, then the flexibility to estimate attribute-specific decision weights may give the IA model the advantage even though it is based on only two attributes out of a larger set of potentially relevant attributes. We also compared a model in which we tried to combine the advantages of the comprehensive overall value ratings together with the flexibility to estimate choice-specific weights for a subset of the individual attributes. We thus fit a third DDM model (IA+) that used the pleasure and nutrition ratings as well as information about the foods’ values beyond those two attributes, in the form of the reported overall value. This provides a measure to quantify the benefit of including individual attribute ratings in the model in addition to overall value ratings. Across all six datasets (studies 0-5 from (Lee & Holyoak, 2021)), the winning model was the IA+, with an exceedance probability of 1 and an estimated model frequency of 0.65. The OV model had an estimated frequency of 0.29, and the IA model had an estimated frequency of 0.06 (Figure 5).

**Figure 5.**
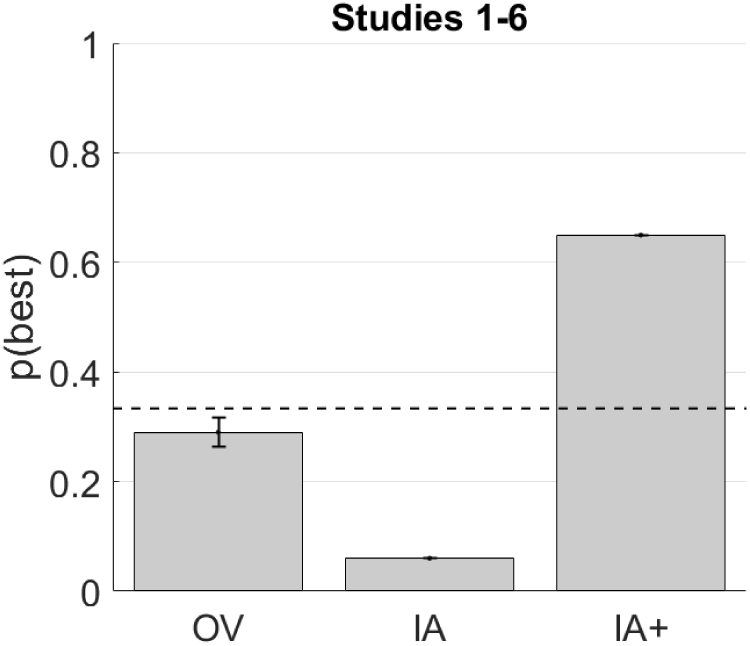
Model comparison results. Simultaneous comparison of the OV, IA, and IA+ models. We show here the probability that each model best explains the data across the participant population, across all studies (n=307). The black dashed line indicates chance level if all models were equally probable a priori.

As a robustness check, we repeated the above model comparisons using the final post-choice ratings from each experiment (i.e., Rating 2 for Experiments 0-1 and Rating 3 for Experiments 2-5). If somehow the initial ratings were too noisy or mis-calibrated, we might expect to find different results when using final ratings to drive the models. The model comparison results using final rather than initial ratings were very similar: IA+ won with an exceedance probability of 1 and an estimated frequency of 0.57, OV had an estimated frequency of 0.35, and IA had an estimated frequency of 0.08. The main difference was that OV gained some support (6%) when using post-choice ratings (consistent with the idea that the accuracy of overall value ratings was refined during choice deliberation; (Lee & Daunizeau, 2020, 2021)).

To test the influence of confidence ratings on the model comparison results reported above (which were based on choice probability, RT, and confidence), we repeated that analysis while excluding confidence data. The assumption that confidence is linearly related to the evidence accumulation rate is arguably the least supported of our three generalized predictions. The model comparison results without confidence were similar to those with confidence, although OV gained a little more support than it did before (exceedance probability: 0.04 for OV, 0.96 for IA+; estimated model frequency: 0.39 for OV, 0.12 for IA, 0.49 for IA+; see also Supplementary results for hierarchical model fits without confidence ratings). This suggests that including confidence data helps to distinguish the models (see model recovery details in the Methods section).

The last model comparison analysis that we performed examined the dataset for which we had overall value ratings as well as four individual attribute ratings (taste, health, texture, appearance). Here, even though this additional dataset had attribute ratings covering a larger range of potentially relevant attributes (4 versus 2), the IA+ version again dominated (exceedance probability = 1; estimated model frequency for IA+ = 0.96, for OV and IA = 0.02 each; but note the small sample size, n=17; Figure 6). To check whether adding additional attribute ratings would always be useful, we compared versions of the IA specification that included only one attribute (taste), two attributes (taste + health), three attributes (taste + health + texture), and four attributes (taste + health + texture + appearance). The version with three attributes performed best (exceedance probability = 0.63, estimated model frequency = 0.42), followed by the version with two attributes (exceedance probability = 0.24, estimated model frequency = 0.31), followed by the version with one attribute (exceedance probability = 0.13, estimated model frequency = 0.26). However, when pitted against OV and the full IA+, the results remained the same as reported above.

**Figure 6.**
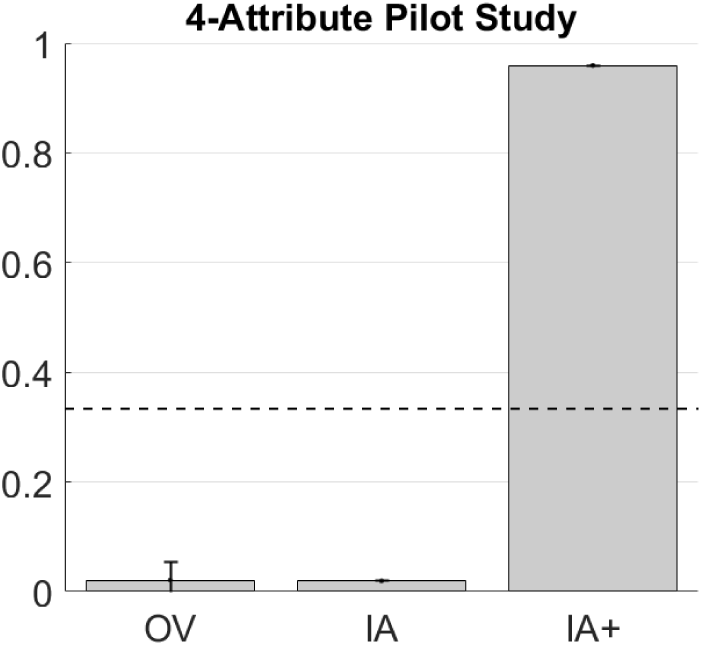
Model comparison results. Simultaneous comparison of the OV, IA, and IA+ models. We show here the probability that each model best explains the data across the participant population, for the four-attribute pilot study (n=17). The black dashed line indicates chance level if all models were equally probable a priori.

In this paper, we do not examine the parameter estimates for the evidence accumulation models for two reasons. First, we are testing general predictions of the evidence accumulation models rather than a specific question about parameter values. Second, we know that overall value is highly correlated with pleasure and nutrition (indeed, it is assumed to be composed of them). This would make it complicated to accurately estimate the different parameters for each rating when fitting the IA+ model to each participant individually, but does not compromise the overall model fit.

## Discussion

Most options that humans need to evaluate and choose from in daily life have multiple attributes or dimensions, and we found that these attributes are combined in a context-dependent manner to determine an option’s value. Previous work on naturalistic multi-attribute decisions has shown that people form option representations based on a large number of separate underlying attributes (Bhatia & Stewart, 2018). Our results on both choice outcomes and post-decision confidence ratings are consistent with these findings. Similar results have also been reported for options where the attributes are objectively defined, such as intertemporal choice (monetary amount and delay (Amasino et al., 2019)) and risky choice (monetary amount and probability (Lee et al., 2022)). In addition, our model simulation and comparison results suggest that the representations that determine an option’s subjective value may differ between rating and binary choice tasks. From the simulation analysis, it is clear that decision models based on a fixed, unitary overall value equal to that given during the ratings sessions do not generate the influence of disparity on choices, response times, or confidence ratings seen in the empirical data (Lee & Holyoak, 2021). A simple explanation for this result is that the influence of pleasure, nutrition, or other attributes differs between the preference elicitation methods (ratings versus choices). In terms of quantitative model comparisons, if the overall values of the foods in these six studies were invariant across the rating and choice tasks, then models using only the overall values to determine the drift rate (OV) would be preferred over models using either a subset of the food attributes (i.e., IA with pleasure and nutrition) or individual attributes plus the overall values (IA+). This is because relative to the OV specification, IA adds complexity (one additional parameter) while at the same time reducing the completeness of the information about the food items – assuming overall value is determined by more than just pleasure and nutrition. Moreover, if overall value representations were constant, then the IA+ specification would add redundant complexity compared to OV and be penalized for that complexity without benefiting from greater explanatory power in the comparisons. In fact, IA+ (overall value + pleasure + nutrition) dominated the other models in the model comparison, as it is the best in terms of generating the observed effects of attribute disparity and accounting for the observed pattern of choice outcomes, response times, and confidence ratings.

We acknowledge the possibility that our model comparison results could be sensitive to the choice of priors for the fitted parameters, which are required input to the VBA procedure that we used. In particular, when we repeated our analyses with certain other reasonable prior means, the model recovery performance dropped substantially. In those cases, the procedure was often unable to distinguish the OV and IA+ models, and so often misattributed true IA+ simulated trials to OV because of the lower complexity of the OV model. Nevertheless, the results we report in the Supplementary Material – based on a more standard model-fitting procedure with typical weakly informative priors – replicate the results we report in the main text, showing that they are robust across model comparison procedures.

The construction of a choice option’s subjective value is an active, malleable process. We have shown that when options are composed of multiple distinct attributes, the manner in which these attributes are evaluated and potentially combined to determine the overall value of each option relative to the other occurs in the moment (i.e., at the time of rating or choice solicitation). Specifically, the contributions of inherent attributes such as pleasure and nutrition to the overall value of food rewards may differ when the foods are evaluated in isolation compared to when choices are made between pairs of foods, even though the goal of the valuation process should be the same in both cases. These findings indicate that preferences over naturalistic multi-attribute goods are labile, and that it might not always be reasonable to assume that previously-obtained overall value ratings summarize the same value representations that guide current choice behavior. Note that our results hold using either pre- or post-choice ratings of overall value, pleasure, and nutrition and regardless of whether the overall value or attribute-specific ratings were given closest to the choice phase, ruling out temporal proximity or item familiarity as potential explanations for these results.

The superior ability of models including attribute-level information to explain the effects of disparity on choices, RTs, and confidence ratings indicates that the importance of one or more attributes reliably differs during binary choices relative to ratings. Specifications of the models that use reports of overall value as the input to the evidence accumulation process implicitly hold the relative importance of each attribute fixed, and thus cannot account for differences in value computation between ratings of single options and choices over two or more options. In contrast, a multi-attribute model specification directly estimates the importance weights for each attribute within the choice context and is therefore better able to explain choice behavior. However, these models are agnostic about how or why the importance of specific attributes differs when individuals are computing the overall value of a single option compared to choosing between two options.

Many sequential sampling models of decision-making posit that attention and salience play an important role in value computation and comparison. Examples of such models are Decision Field Theory (DFT; (Busemeyer & Townsend, 1993)), an extension of DFT known as the Multi-Attribute Dynamic Decision (MADD) model (Diederich, 1997), and the attentional DDM (aDDM; (Krajbich et al., 2010; Smith & Krajbich, 2019)). In the DFT model, the drift rate can vary across deliberation time if, for example, one option is more salient but the other is truly more valuable. The MADD model makes the multi-attribute nature explicit, and the drift rate fluctuates over time as the decision maker shifts focus across the set of relevant attributes. Although the aDDM has generally been applied to the overall values of options, or to distinct items within a bundle (Fisher, 2017, 2021b), it would be conceptually similar to the MADD if applied at the attribute level for goods that are inherently multidimensional. Query theory is similar to these sequential sampling models in that it holds that the order in which a decision maker considers different aspects (or attributes) of an option alters its resultant valuation (E. Johnson et al., 2007; Weber et al., 2007). In all these models, it is assumed that options or attributes that receive more attention will be favored during the value comparison process. However, the relationship between value and attention is most likely bidirectional to some extent (Anderson et al., 2011; Callaway et al., 2021; Gluth et al., 2018; Jang et al., 2021; Towal et al., 2013; Westbrook et al., 2020).

Differences in the amount of attention directed to specific attributes during the evaluation and decision contexts could explain changes in the manner in which the attributes aggregate into overall value across those contexts. Consistent with this idea, changes in the proportions of visual fixations to locations on a computer screen indicating the monetary amount versus the probability of winning when pricing versus choosing between lotteries are associated with inconsistencies between the two preference elicitation contexts (Alós-Ferrer et al., 2021; Kim et al., 2012).

Fixation patterns towards monetary amount versus delay affect temporal discounting rates (Fisher, 2021a), although whether this might vary in pricing versus choice contexts has not been tested. Covert attention processes may have effects similar to visual attention for naturalistic multi-attribute goods. The focus of covert attention is more difficult to measure than visual attention (i.e., fixation locations), but studies combining decision tasks with neuroimaging or electrophysiology and machine learning techniques may give us a window into these cognitive processes (Aoi et al., 2020; Peixoto et al., 2021). Experimental manipulation of focus on a specific attribute (Fisher, 2018; Hare et al., 2011) may also prove useful, if an appropriate method to dynamically shift attention within each trial is developed.

In addition to providing further insight into the mechanistic nature of value-based decisions, our current work has practical implications for future studies of decision-making. We have shown that it is best to use as much attribute-level information as possible when modeling decisions over multi-attribute stimuli. Most, if not all, naturalistic stimuli are composed of multiple attributes, thus most studies of decision-making should incorporate attribute-level information. At the same time, it will often be impractical to collect information on a large number of attributes, especially if one needs subjective opinions about the attributes from each participant in an experiment (but see the Supplementary Material regarding using population-level averages as proxies for additional attribute ratings). Our results indicate that combining attribute-specific and overall values may be a good compromise between attempting to include comprehensive attribute-level information and conforming to practical constraints. Naturally, which attribute-level information to obtain and how to best combine it with some type of overall value rating will depend on the hypotheses and experimental design. Given the clear evidence that the value-comparison process is based on context-dependent attribute assessments, experiments that use a well-designed combination of attribute-specific and aggregate-level information should prove to be the most useful in advancing our understanding of many important decision mechanisms. We believe that the datasets we examined in this study may be the only ones currently available that include subjective ratings of both overall value and individual attributes. It would be beneficial if future studies also collected such comprehensive data, for without it, effects such as those we have shown here might never be exposed.

## Data and Analysis Code Availability

The raw data and primary analysis code for this study are available in the Open Science Forum at http://doi.org/10.17605/OSF.IO/EDSH6.

## Supplementary materials

### Correlation between pleasure and nutrition ratings

In this study, we examined the relative subjective importance of the pleasure (P) and nutrition (N) attributes when people either rate individual snack foods or choose between pairs of snack foods. It is thus beneficial if ratings along these two attribute dimensions are not highly correlated, as that would make the regression analysis results less straightforward to interpret. In the main text, we reported that the average Pearson correlation between P and N (across participants) was not statistically different from zero. However, we did find a good deal of heterogeneity across participants, such that some provided P and N ratings that were negatively correlated, while others provided P and N ratings that were positively correlated. Figure S1 displays a histogram of the correlation between P and N for the entire participant pool. Note that the majority of participants (about 80%) had correlation coefficients between -0.5 and 0.5.

**Figure S1:**
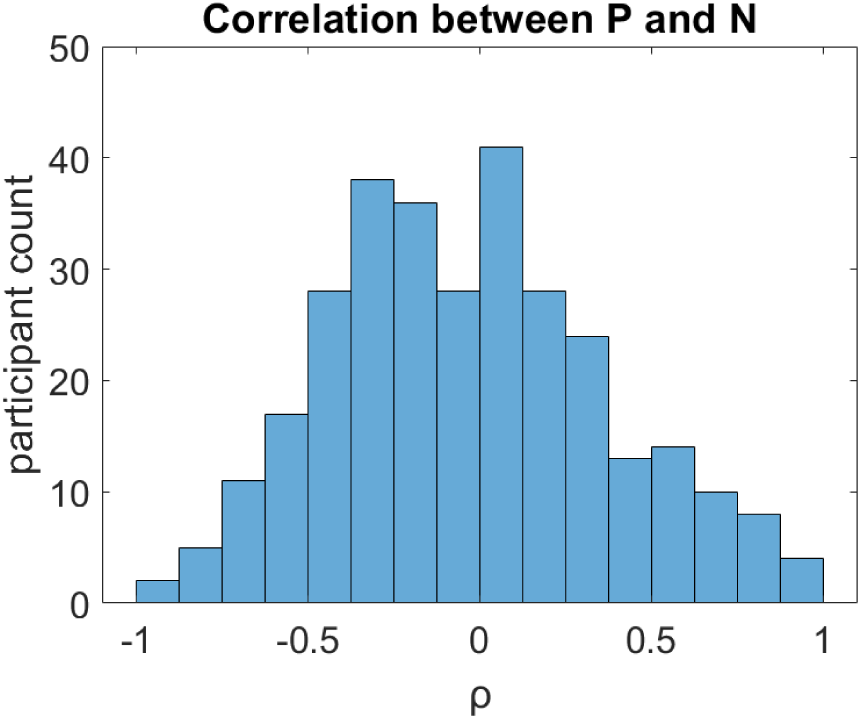
Histogram of participants’ coefficients of correlation between P and N.

### Correlation between predicted and empirical variables

As an additional check that our models did a reasonable job of accounting for the empirical data, we compared the predicted RT and confidence ratings (under each model formulation) with the true empirical data. As shown in Figure S2, each model did a good job of accurately predicting the data (all correlation levels are around 60%). Note that the correlations between data simulated under the IA+ formulation show the highest correlation with the empirical data, which is to be expected given that the IA+ outperformed the other formulations in the formal model comparison.

**Figure S2:**
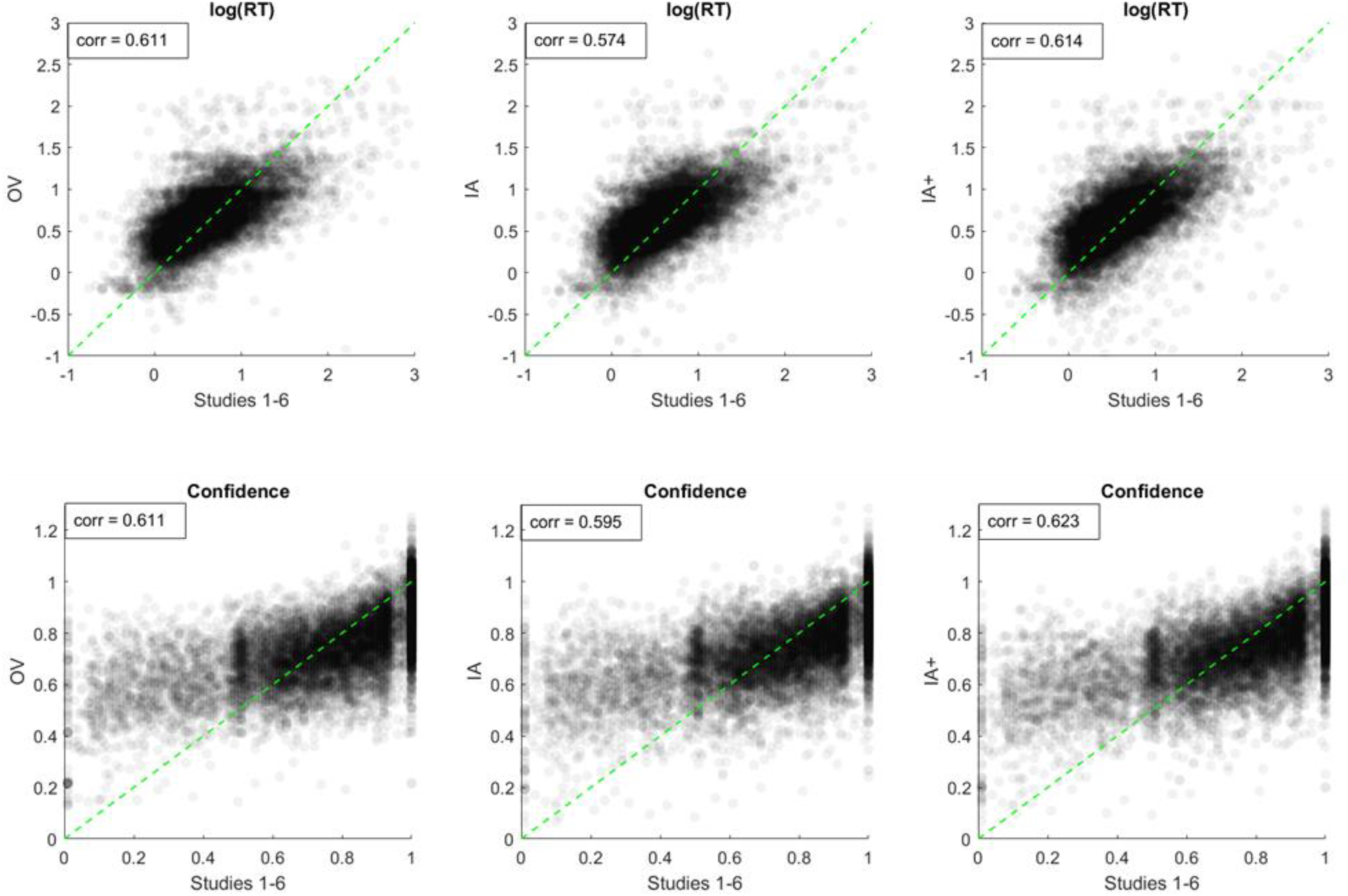
Correlations between data (RT and confidence) predicted under each model formulation and empirical data. The upper row shows the associations between the logarithms of empirical response times (x-axis) and response times simulated under each evidence accumulation assumption (y-axis). The lower row shows the associations between empirical confidence ratings (x-axis) and confidence ratings simulated under each evidence accumulation assumption (y-axis). The green dotted lines show the identity line for all plots.

### Consistency of attribute ratings across participants

In the main text, we demonstrated that it is beneficial to include ratings of multiple individual attributes when modeling choice behavior, in addition to the traditional ratings of overall value. However, it might not always be practical to collect ratings for many different attributes (e.g., due to time constraints or participant boredom or fatigue). As a potential solution for this dilemma, one might wonder whether individual subjective ratings could be replaced (as an approximation) by population-level averages. If true, an experimenter could accumulate a database of ratings of many attributes for many options, because each attribute could in principle be evaluated by a different group of participants (and used to model the behavior of other participants in future studies). In order for such a method to work, there would have to be a high degree of similarity in the ratings for each option across participants. Whereas this might be unlikely for ratings based largely on personal experience and personal preferences (e.g., value or pleasure), it might be reasonable for ratings based more on learned information (e.g., nutrition), especially when such learning is generally consistent across a specific culture (e.g., for the residents of the US and Canada in this study). We thus calculated aggregate statistics (mean and standard deviation) for the 100 items in Study 2 (Study 1 did not include an identical set of items across participants) and for the 60 items in Studies 3-6 across all participants in those studies. As shown in Figure S3, there is a lot of variability in the value and pleasure ratings, but much less variability in the nutrition ratings.

**Figure S3:**
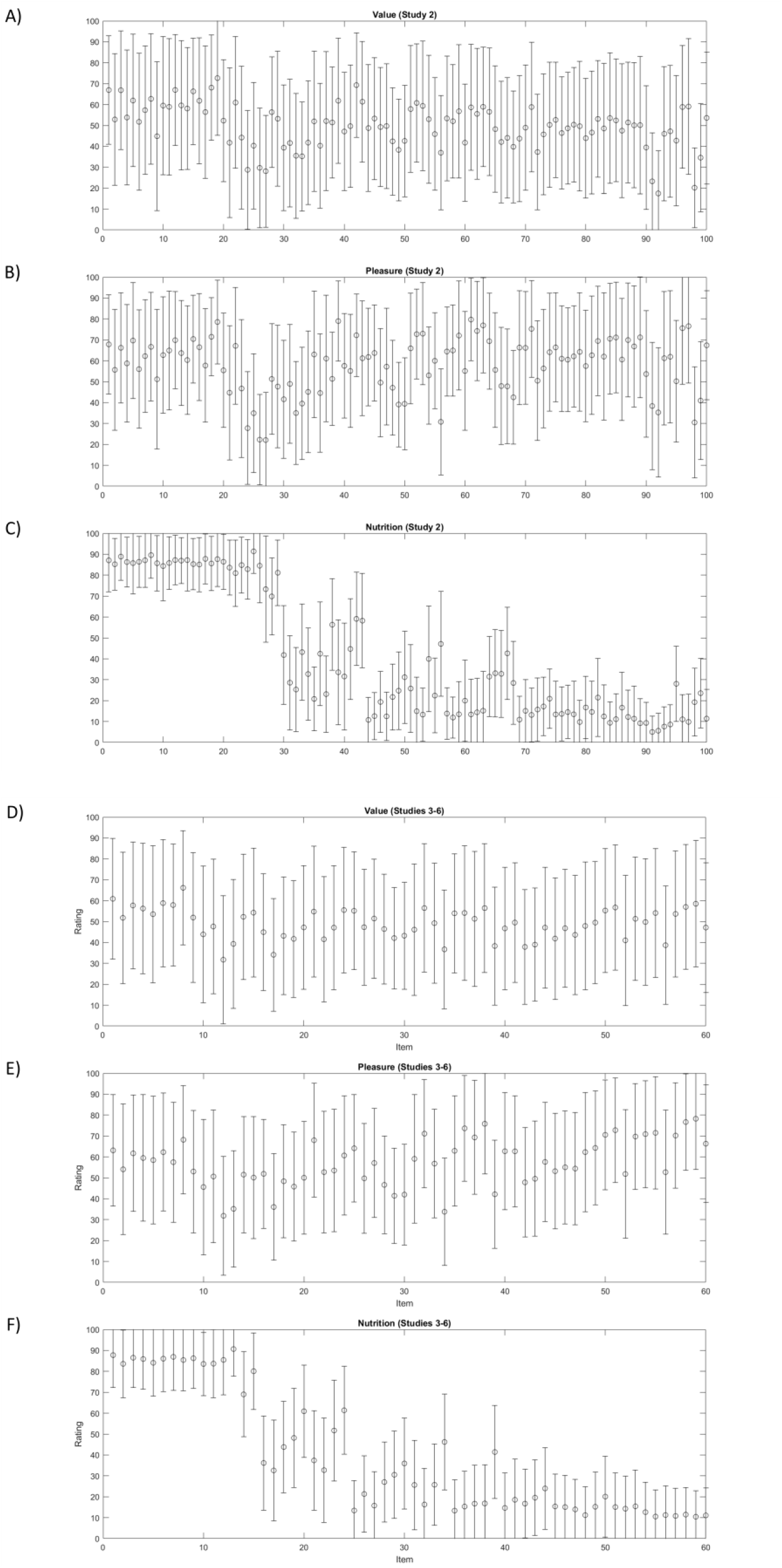
Ratings across participant populations. Population statistics for the ratings of value (panels A & D), pleasure (panels B & E), and nutrition (panels C & F) for participants in Study 2 (panels A-C) and Studies 3-6 (panels D-F). Circles indicate means, error bars indicate +/- one standard deviation.

### DDMs with log-likelihoods based on the full distribution of predicted RTs

#### Fitting methodology

In addition to the simplified fits using mean RTs, consistency, and confidence reported in the main text, we also fit the overall value (OV), individual attribute (IA), and individual attribute plus overall value (IA+) models using the entire response time distributions predicted by the drift diffusion model (DDM) to compute the log-likelihood of proposed parameter combinations with the R package runjags (Denwood, 2016; R Core Team, 2020) as an interface for JAGS (Plummer, 2003; Wabersich & Vandekerckhove, 2014). For all three DDMs, we fit hierarchical Bayesian models using three parallel Markov Chain Monte Carlo samplers (50,000 burnin samples; 10,000 posterior samples with a thinning step = 1 per chain). We used uniform priors for the drift noise (0.001, 1) and non-decision time (0, 3), and standard normal distributions (mean = 0, sd = 1) for the 1-3 drift scaling parameters in each model (*d^o^*, *d^a^*, and *d^b^*, see main text methods for further details). We fixed the boundary separation value at 2. Lastly, initial fits to all six data set showed that there was no bias to select the left or right item, and therefore we fixed the bias parameter at 0.5 (no bias) when fitting data from studies 1-3 and 4-6 separately to test the out-of-sample log-likelihoods for the OV, IA, and IA+ models. The hierarchical model allowed for participant-specific drift scaling and noise parameters as well as non-decision times. For all fits, we excluded trials with response times that were less than 0.2 seconds or greater than 10 seconds (less than 0.5% of the data). The plot shown in Figure S4 was created using the R packages ggplot and GGally (Schloerke et al., 2021; Wickham et al., 2022).

### Model Comparison

We fit each of the three DDMs to the data from studies 1-3 and 4-6 separately and then computed the log-likelihood of the data observed in the three left out studies to compare the out-of-sample performance of the DDMs. The out-of-sample log-likelihoods (summed over both tests) of the OV, IA, and IA+ were -20742, -22610, and -19327, respectively (log-likelihoods closer to 0 indicate better predictions for the new data). Thus, the relative ranking of the three models agrees when fitting them using either the mean or entire RT distributions predicted by the DDM. The IA+ model, which specifies the drift rate as a function of individual attributes and the overall value is the best in both tests.

### Parameter recovery

We tested the accuracy of parameter recovery from the best-fitting IA+ model. Note that the other two models are special cases nested within the IA+ model. In the OV model, the drift scaling parameters on pleasure and nutrition are fixed to zero. While in the IA model, the drift scaling parameter on overall value is fixed to zero. The tables below list: the mean and standard deviations of the group-level parameters from the IA+ model fit to empirical data (left two columns), and corresponding values for fits to data generated using an IA+ DDM with the best-fitting individual-level parameters from the empirical fits (Figure S4; Table S1), and the correlations between generating and recovered individual-level parameters (Table S2).

**Table S1.** Group-level parameter recovery.

|  | **Generating** |  | **Recovered** |  |
| --- | --- | --- | --- | --- |
| **Parameter** | **Mean** | **SD** | **Mean** | **SD** |
| Nutrition | 0.212 | 0.050 | 0.197 | 0.044 |
| Pleasure | 1.120 | 0.085 | 1.054 | 0.075 |
| Overall value | 1.547 | 0.095 | 1.573 | 0.091 |
| Non-decision time | 0.875 | 0.022 | 0.875 | 0.021 |
| Drift noise | 0.827 | 0.016 | 0.828 | 0.012 |

**Table S2.** Correlations between individual-level generating and recovered parameters.

|  |  |  |  | Non- |  |
| --- | --- | --- | --- | --- | --- |
|  | Nutrition | Pleasure | Overall<br>value | decision<br>time | Drift noise |
| Nutrition | 0.680 | -0.025 | 0.180 | -0.025 | 0.154 |
| Pleasure | -0.025 | 0.779 | 0.529 | 0.043 | 0.388 |
| Overall value | 0.162 | 0.462 | 0.831 | 0.097 | 0.392 |
| Non-decision time | -0.092 | 0.097 | 0.083 | 1.000 | -0.268 |
| Drift noise | 0.192 | 0.403 | 0.381 | -0.268 | 1.000 |

951 Values represent Pearson correlation coefficients.

**Figure S4.**
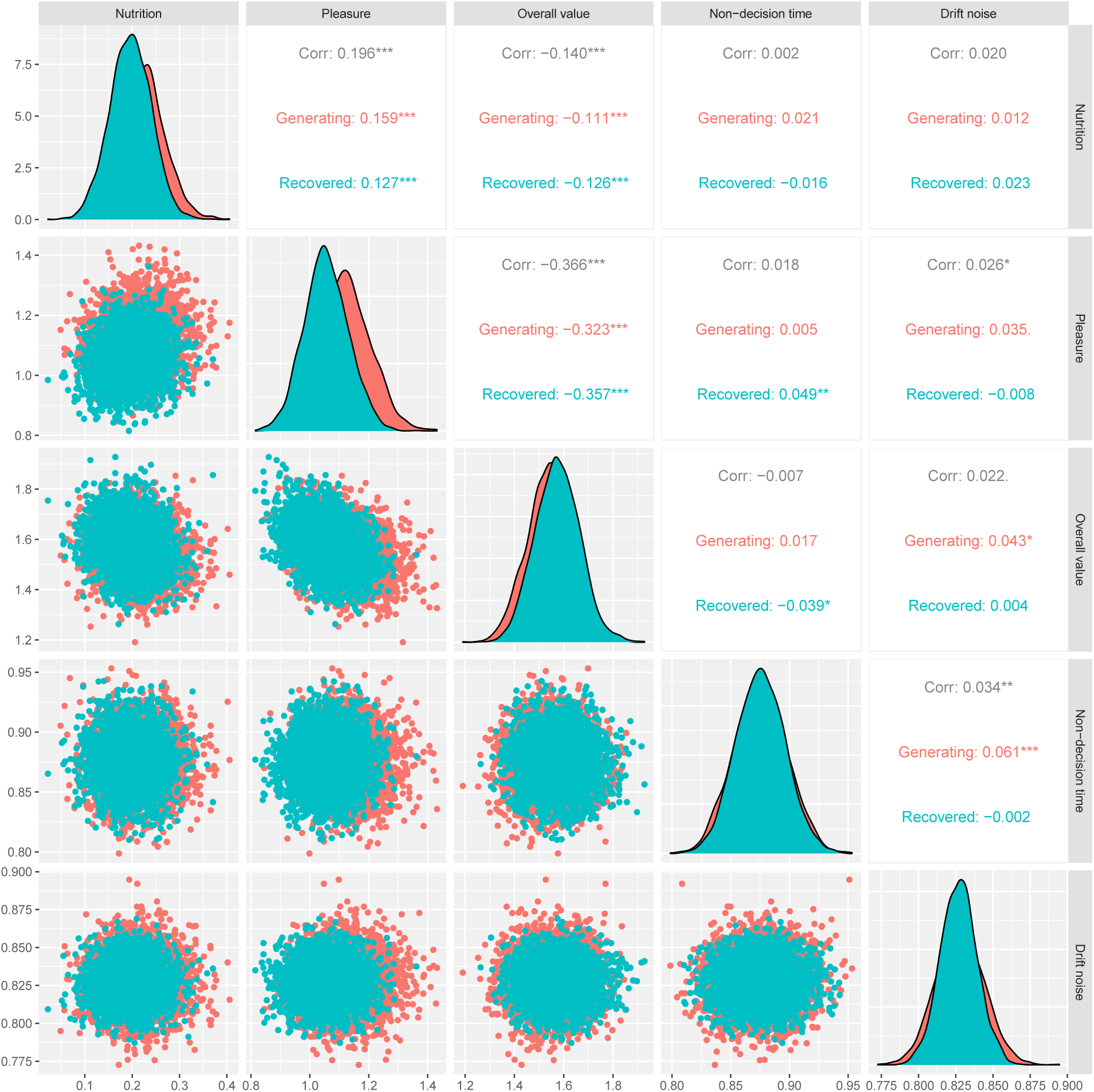
Group-level parameter posterior distributions for the IA+ model fit to the empirical data (Generating, red) and data simulated using the fitted individual-level empirical parameters to generate choices and response times based on the set of trials the participants completed. The density plots along the diagonal show the posterior distribution for each parameter in both fits. The off-diagonal cells show scatterplots of the correlations between posterior chains in each fit (lower) and the Pearson correlation values (upper).

## Footnotes

1 This simple calculation prevents the sampled parameter estimates from ever being negative, while simultaneously preventing them from being overly positively-skewed. Importantly, it does not impede the VBA model-fitting algorithm. We thank Jules Brochard for this tip.

